# Effects of ageing and frailty on circulating monocyte and dendritic cell subsets

**DOI:** 10.1101/2023.08.07.552240

**Authors:** Rosanne D. Reitsema, Ashok K. Kumawat, Bernd-Cornèl Hesselink, Debbie van Baarle, Yannick van Sleen

## Abstract

Ageing is associated with dysregulated immune responses, resulting in impaired resilience against infections and a low-grade inflammation known as inflammageing. Frailty is a measurable condition in older adults characterized by decreased health and physical impairment. Dendritic cells (DCs) and monocytes play a crucial role in initiating and steering immune responses. To assess whether their frequencies and phenotypes in the blood are affected by ageing or frailty, we performed a flow cytometry study (14 markers) on monocyte and DC subsets in an immune ageing cohort (SENEX).

Participants were divided into three groups (n=15 each): healthy young controls (HYC, median age 29 years), healthy older controls (HOC, 73 years) and Frail older controls (76 years). Frailty status was based on Tilburg Frailty Index scores. Among HLA-DR+/CD19-cells, monocyte subsets (classical, intermediate, non-classical) were identified by CD14 and CD16 expression, and DC subsets (conventional (c)DC1, cDC2, plasmacytoid (p)DC) by CD11c, CD1c, CD141 and CD303 expression.

Using unsupervised and conventional gating strategies, we observed a substantially lower proportion of pDCs in the HOC compared to the HYC. Additionally, we observed higher expression of activation markers on classical and intermediate monocytes and on cDC2 in HOC compared to HYC. Comparing the Frail to the age-matched HOC group, we observed only one important difference: a higher expression of CD40 on classical and non-classical monocytes in Frail individuals.

In this cross-sectional study, we document a substantial effect of ageing on monocytes and DCs. The reduction of pDCs in older people may underly their impaired ability to counter viral infections, whereas the enhanced expression of activation markers could indicate a state of inflammageing. Future studies could elucidate the functional consequences of CD40 upregulation with frailty.

## INTRODUCTION

There are major differences in how individuals and their immune systems respond to ageing. The aged immune system is characterized by a dysregulated adaptive immune response to pathogens. On the other hand older individuals often demonstrate signs of enhanced innate immune responses, characterized by the production of pro-inflammatory cytokines leading to low-grade inflammation, referred to as inflammageing. Inflammageing appears to go hand in hand with a shift of the leukocyte composition towards the myeloid lineage, a process also observed in autoinflammatory diseases (1,2).

Frailty is a term used to describe older adults with a decreased health leading to physical impairment, morbidity and is strongly associated with mortality (3,4). It describes the lack in ability of a person to cope with external stressors and results in an inability to handle essential daily tasks (5). The frail population is estimated to comprise between 4% and 59% of the older population. This wide range of estimates is caused by a lack of standardized measurements, different age cohorts and different populations (6). Multiple different approaches have been employed to quantify frailty. These include the Frailty Phenotype (7), the Frailty Index (8), the Groningen Frailty Indicator (9) and the Tilburg Frailty Indicator (TFI) (10) amongst others. These questionnaires and measurements focus on independence at undertaking various activities, physical health, mental health, social and cognitive components. Due to ageing of the worldwide population, frailty is an increasingly important topic (6).

Proper functioning of antigen presenting cells (APCs) is essential for initiating and fine-tuning immune responses. One of the main functions of APCs is to patrol their environment to detect danger signals via pattern recognition receptors (PRRs) (11). Circulating APCs include subsets of dendritic cell (DCs) and monocytes, that are characterized by high expression of HLA-DR, allowing them to present processed antigens to T- and B-cells. In the blood, DCs have been described as ‘immature’, but after activation they typically migrate to the secondary lymphoid organs to initiate the adaptive immune response. Subsets of DCs substantially differ in phenotype and function, and comprise the rare CD141+ conventional DCs (cDC1), the CD1c+ conventional DCs (cDC2), and the CD303+ plasmacytoid DCs. Monocytes may be less specialized in antigen presentation, and are rather known for their phagocytic capability and their cytokine production. The majority of monocytes are classical CD14^+^CD16^-^ monocytes, of which some may develop into CD16^+^ intermediate and non-classical monocytes (12,13).

Considering the importance of APCs in shaping the immune system in ageing individuals, we investigated whether their frequencies and phenotypes in the blood are affected by ageing or frailty. Previously, elevated frequencies of myeloid cells have been associated with frailty (14). However, less is known about specific subsets and their extended phenotype. Therefore, the goal of this study is to document APC subset proportions and phenotypes, in healthy young and elderly individuals, and to compare these with individuals deemed as frail, based on the TFI. To investigate this, we employed an extensive flow cytometry panel to analyse both DC and monocyte subsets.

## METHODS

### The SENEX cohort

APCs were investigated in peripheral blood mononuclear cell (PBMC) samples of participants of the SENEX cohort in the University Medical Center Groningen (UMCG), which was initiated in 2011 to investigate immune ageing in young and older adults. Exclusion criteria were: pregnancy, clinical signs of severe anemia, disease that influence the immune system ((auto)immune diseases, active infection, malignancy) and drugs that influence the immune system (corticosteroids, recent vaccination). In this study, we included 15 younger participants (HYC, <35 years), and non-frail (HOC) and frail older participants (>50 years, n=15 each). All participants signed for informed consent and all procedures were in compliance with the declaration of Helsinki. The study was approved by the institutional review board of the UMCG (METc2012/375).

### Frailty assessment

The older participants in the SENEX cohort completed a number of questionnaires on health and frailty at each visit. The defining criterium for frailty was based on the TFI scores (10), which ranged from 0-15 points. The TFI consists of three components, physical, psychological and social. Participants in our Frail group had to have a score of ≥5, or a score of 4 with a score of ≥2 on the physical component. Non-frail participants had a score of ≤1 on the TFI. Additionally, participants were scored on the SF-36, GFI and HAQ questionnaires, and the Fried Frailty Phenotype (7,9,15,16).

### Flow cytometry

APCs were investigated using a 14-color flow cytometry panel on thawed PBMCs (Supplementary Table 1). Cells were measured on the BD FACSymphony flow cytometer, whose laser settings were normalized every day prior to measurement using cytometer setup and tracking beads. Fluorescent compensation was done initially with compensation beads and finalized using fluorescence minus one controls. To quantify the proportions and phenotypic expression of the APCs, manual gating analysis (Supplementary figure 1) was performed with Kaluza V2.1 software (Beckman coulter, IN, USA).

**Table 1:** Participant characteristics of the three study populations. Values are given as median and total range. Laboratory data were measured on the XN-9000 (Sysmex, Kobe, Japan), including the diff values for monocytes and PBMCs (lymphocytes+monocytes). Statistically significant differences, tested by the Mann Whitney U test, of the Frail or HYC groups compared to the HOC group are indicated by a ‘*’. HOC: Healthy control, HYC: Healthy young control, ESR: erythrocyte sedimentation rate, Hb: haemoglobin, PBMCs: peripheral blood mononuclear cells, TFI: Tilburg Frailty Indicator, GFI: Groningen Frailty Indicator, SF-36: Short Form 36, HAQ: Health Assessment Questionnaire.

|  | HOC | Frail | HYC |
| --- | --- | --- | --- |
| **n** | 15 | 15 | 15 |
| **Age (years)** | 73 (66-82) | 76 (59-95) | 29 (25-32) |
| **Sex (male/female)** | 6/9 | 4/11 | 8/7 |
| **Laboratory data** |  |  |  |
| **ESR mm/h** | 10 (3-28) | 10 (1-34) | 4 (1-15) * |
| **Hb mmol/L** | 9.0 (7.6-10.2) | 8.7 (7.2-9.7) | 9.2 (7.8-10.3) |
| **PBMCs (10<sup>9</sup>/L)** | 2.5 (1.5-3.6) | 2.1 (1.4-3.8) | 2.6 (1.5-3.3) |
| **Monocytes (10<sup>9</sup>/L)** | 0.56 (0.24-0.82) | 0.55 (0.22-0.73) | 0.41 (0.21-0.64) * |
| **TFI** | *n*=15 | *n*=15 |  |
| **Total score** | 0 (0-1) | 6 (4-11) * |  |
| **Physical score** | 0 (0-1) | 3 (1-7) * |  |
| **Psychological score** | 0 (0-1) | 3 (0-3) * |  |
| **Social score** | 0 (0-1) | 2 (0-3) * |  |
| **Fried frailty phenotype** | *n*=15 | *n*=10 |  |
|  | 0 (0-0) | 0.5 (0-4) * |  |
| **GFI** | *n*=15 | *n*=15 |  |
| **Total score** | 0 (0-3) | 4 (0-9) * |  |
| **Physical score** | 0 (0-1) | 0 (0-4) |  |
| **Mental score** | 0 (0-3) | 3 (0-6) * |  |
| **SF-36** |  |  |  |
| **Physical functioning** | 95 (70-100) | 75 (40-100) * |  |
| **Role of physical limitations** | 100 (50-100) | 100 (0-100) |  |
| **Role of emotional limitations** | 100 (100-100) | 67 (0-100) * |  |
| **Energy/fatigue** | 85 (70-100) | 65 (35-90) * |  |
| **Emotional well-being** | 84 (68-100) | 64 (32-92) * |  |
| **Social functioning** | 100 (62.5-100) | 75 (50-100) * |  |
| **Pain** | 100 (45-100) | 67.5 (25-100) * |  |
| **General health** | 70 (55-95) | 60 (35-95) * |  |
| **HAQ damage index** | *n*=9 | *n*=13 |  |
|  | 0 (0-0.56) | 0.28 (0-2) * |  |

**Figure 1:**
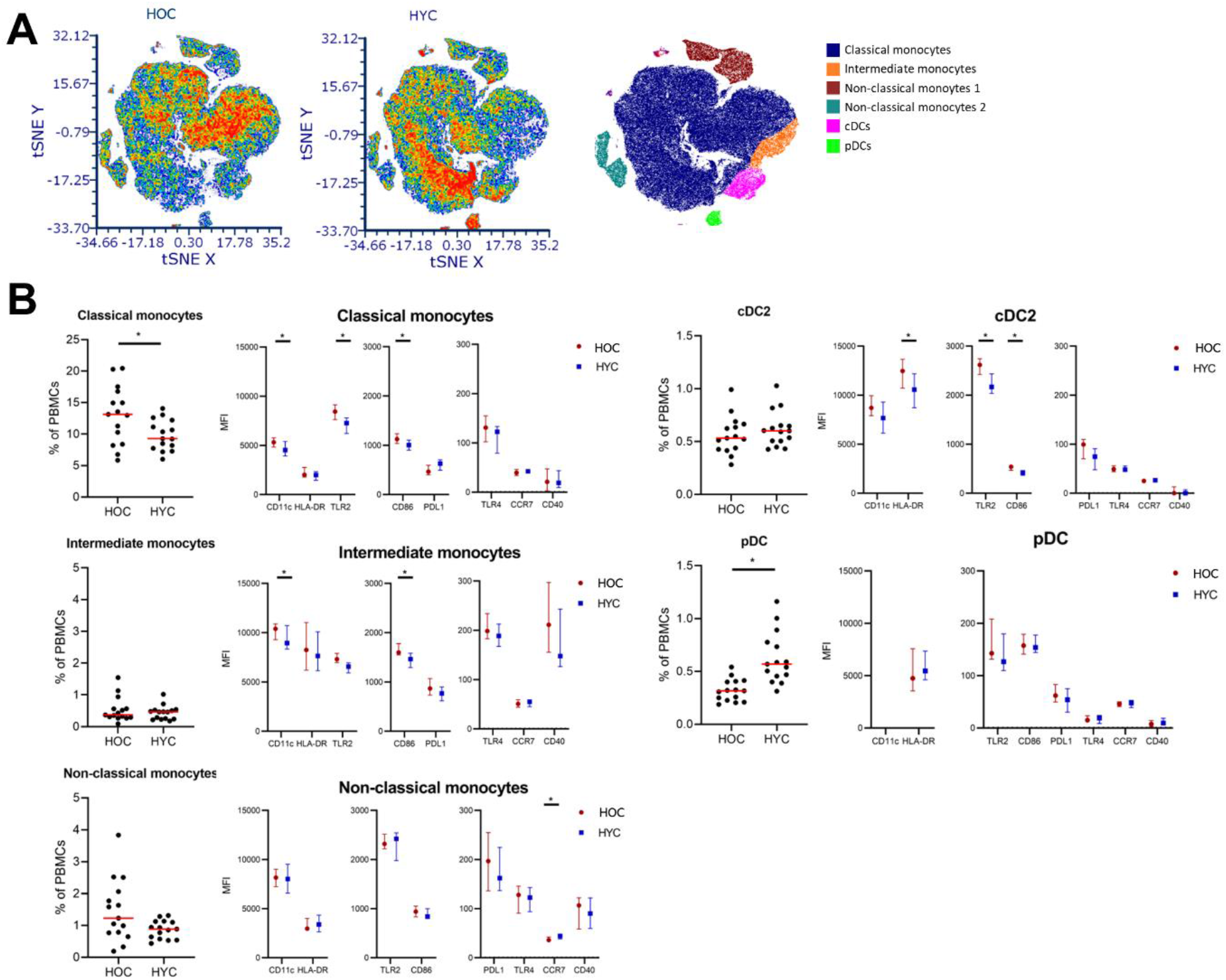
Changes in APC frequencies and phenotypes with age. Shown in A are t-distributed stochastic neighbor embedding (t-SNE) plots of HOC (n=15) and HYC (n=15), including a plot indicating the localization of clusters with classical monocytes (CD14+CD16-), intermediate monocytes (CD16+CD14+), non-classical monocytes (CD14lowCD16+), conventional dendritic cells (CD141/CD1c+) and plasmacytoid dendritic cells (CD303+). In B, we show the proportion within total PBMCs of the APC subsets between the HOC and HYC groups. The red line indicates the median and the p-values of Mann Whitney U testing are indicated in the graph. Additionally, we show the MFI for eight markers for each subset, for the HOC and HYC groups. These data are expressed as median +/-inter-quartile range, and statistical differences (P<0.05) in expression between the groups, by Mann Whitney U, is indicated with ‘*’. APC: antigen-presenting cell, HOC: healthy control, HYC: healthy young control, PBMC: peripheral blood mononuclear cells, MFI: mean fluorescence intensity.

### Statistics

To compare HYCs and HOCs, and the Frail and HOC groups, Mann-Whitney U tests were performed, as data were not normally distributed. P-values <0.05 were considered statistically significant. Graphs and statistical tests were created in GraphPad Prism V9. Additionally, we performed t-distributed Stochastic Neighbor Embedding (t-SNE) analyses using FCS express version 6 (De Novo software, CA, USA). T-SNE was calculated on all APCs gated as HLA-DR+CD19-cells merged into one file including a file identifier. The calculation was based on expression of CD14, CD16, CD303, CD1c, CD141, PDL1, CD40, CD86, TLR2, HLA-DR and

CD11c. Sampling options included an interval down sampling method, a Barnes-Hut approximation of 0,50, perplexity set to 60 and number of iterations to 1800.

## RESULTS

### The effects of ageing on the frequencies and phenotypes of monocyte and DC subsets

We first performed t-SNE analysis to help visualize the high-dimensional data onto a two-dimensional plot. Several clusters could be observed after transforming the flow cytometry data using t-SNE. The distribution of APCs appeared to differ between younger and older individuals (Figure 1A).

After assigning cell subsets to each cluster using the expression of each lineage marker (Figure 1A, Supplementary figure 2) it appears that the main distribution and phenotype differences were observed within the classical monocytes cluster (CD14+, CD16dim). Furthermore, the population of pDCs (CD303+) seemed to be much smaller in HOC than in HYC. Using conventional gating strategies (Supplementary figure 1) we further analyzed the distribution and phenotype of each APC subset to assess whether this changed upon ageing.

The main difference in the proportions of APCs observed between younger and older healthy adults is indeed a substantially lower proportion of pDCs in older individuals (p<0.001, Figure 1B). No differences in the proportions of the cDC1 (Supplementary figure 3) and cDC2 subsets (Figure 1B) were observed. Monocyte counts in the blood were higher in the older group (Table 1), although this did not appear to depend on one specific subset, as proportions of both classical and non-classical monocyte subsets within total PBMCs tended to be higher in the HOC group. No shifts with age were observed for monocyte subsets as a percentage within total monocytes (Supplementary Figure 4).

The expression of a number of activation markers on certain APC subsets changed with age (Figure 1B). The expression of pattern recognition receptor TLR2, co-stimulatory signaling molecule CD86, and cell adhesion molecule CD11c was higher on classical monocytes of HOCs than of HYCs. This elevated expression was also found for CD86 and CD11c on intermediate monocytes. Similarly, cDC2 cells of HOCs also expressed higher levels of TLR2, CD86 and also HLA-DR than those of HYCs. CCR7 expression was very low on all subsets but we observed a significantly higher expression on non-classical monocytes of younger compared to older participants. No differences in expression were observed for pDCs, and expression of the markers on the cDC1 subset was not assessed due to their very low frequency.

### APC frequencies and phenotypes in frail and non-frail people

Next we compared the HOC group with the age-matched Frail group, whose frailty status was based on the TFI. Complementary to the TFI, we also observed substantial differences in other quality of life outcomes between the HOC and Frail groups, but no significant differences in general laboratory markers such as ESR and monocyte counts (Table 1).

The tSNE analyses show distribution differences between HOC and Frail within the classical monocyte population, but no apparent differences in other subsets (Figure 2A, Supplementary figure 5). Our subsequent analyses of the monocyte and dendritic cell subset proportions did not show any significant differences between the HOC and Frail groups (Figure 2B,

**Figure 2:**
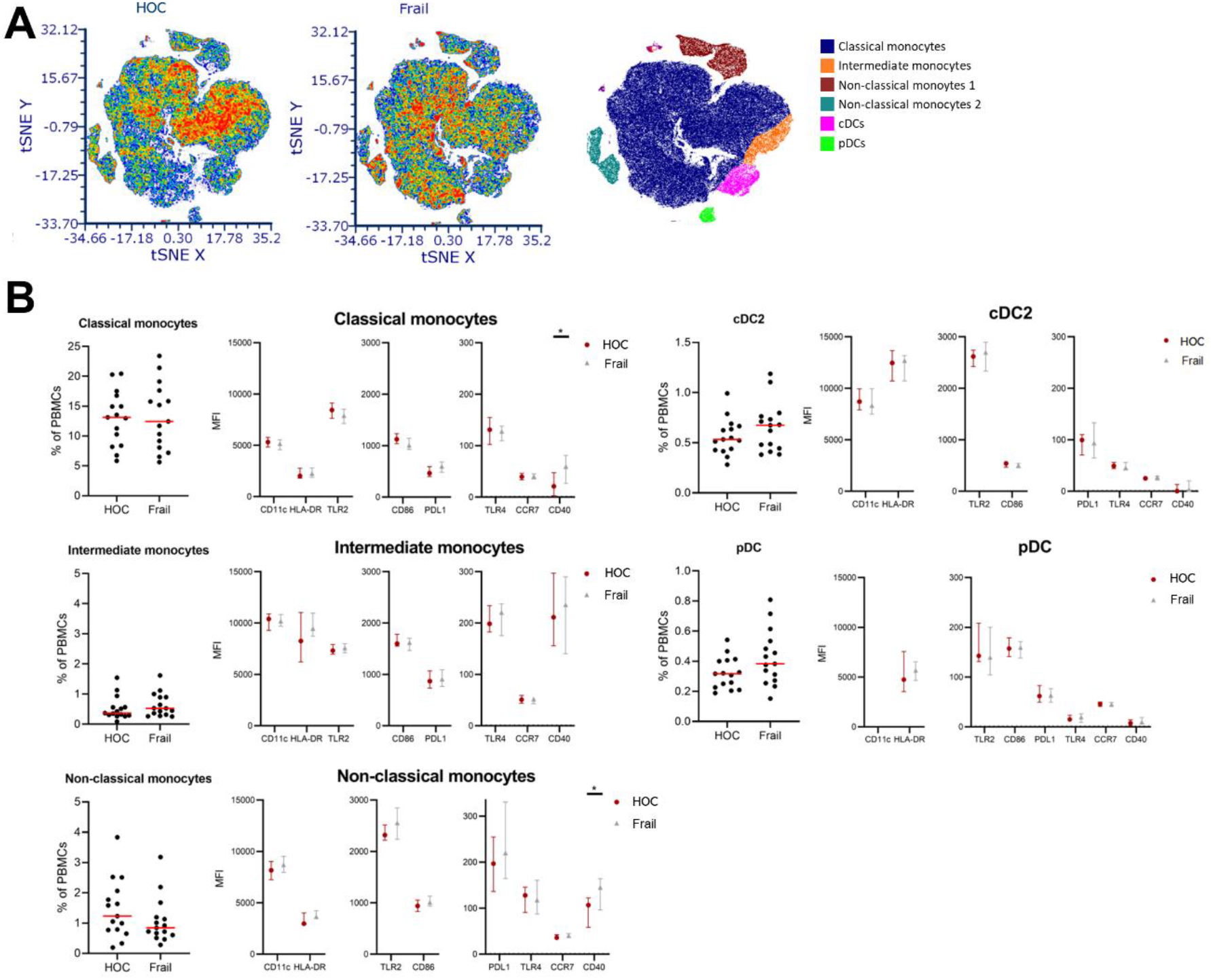
A comparison in APC frequencies and phenotypes between age-matched healthy and frail older adults. Shown in A are t-distributed stochastic neighbor embedding (t-SNE) plots of HOC (n=15) and Frail older adults (n=15), including a plot indicating the localization of clusters with classical monocytes (CD14+CD16-), intermediate monocytes (CD16+CD14+), non-classical monocytes (CD14lowCD16+), conventional dendritic cells (CD141/CD1c+) and plasmacytoid dendritic cells (CD303+). In B, we show the proportion within total PBMCs of the APC subsets between the HOC and Frail groups. The red line indicates the median and the p-values of Mann Whitney U testing are indicated in the graph. We also show the MFI for eight markers for each subset, for the HOC and Frail groups. Data are expressed as median +/-inter-quartile range, and statistical differences (P<0.05) in expression between the groups, by Mann Whitney U, is indicated with ‘*’. APC: antigen-presenting cell, HOC: healthy control, PBMC: peripheral blood mononuclear cells, MFI: mean fluorescence intensity.

Supplementary figure 6). We did not observe any differences between frail and non-frail people in the expression of markers that were shown to increase with age (e.g. CD86, CD11c). CD40, which is essential in APC-T cell communication, was higher expressed in the Frail group compared to the HOCs. CD40 expression on classical monocytes was practically non-existent in the HOC group but was clearly expressed in most frail people (p=0.02). The expression of CD40 on non-classical monocytes is higher than on classical monocytes but was particularly high on non-classical monocytes in the Frail group (p=0.02). Additionally, we observed two trends, with lower CD86 expression on classical monocytes and higher PDL1 expression on non-classical monocytes of Frail people.

## Discussion

In summary, our study explored the effects of ageing and frailty on the frequencies and phenotypes of monocyte and dendritic cell (DC) subsets. Through comprehensive analyses, we identified significant differences in the proportions of pDCs between younger and older individuals, indicating a substantial decline in pDCs with age. However, age-related changes in the expression of activation markers were more prominent on monocyte subsets and cDC2s. Furthermore, comparing a group of healthy and frail older individuals, we found no discernible differences in APC frequencies or phenotypes, except for higher expression of CD40 on classical and non-classical monocytes among the frail group. These findings provide valuable insights into the impact of ageing on APC populations and highlight potential implications for immune function in older adults.

The increased expression of TLR2, CD86, and CD11c on monocytes and HLA-DR, TLR2 and CD86 on cDC2s with age indicates a heightened activation status of these cells. A heightened activation status of cDC2 cells could be reflective of increased maturation, which was previously found to be related to ageing in a study on cDC2 maturation in several lymphoid tissues (17). Increased maturation of cDC2 cells could lead to more readily stimulation of naïve T cells. We only observed higher HLA-DR expression by cDC2 upon ageing, as opposed to other studies which have previously found higher expression of HLA-DR by monocytes as well, along with some other adhesion and migration markers (CD11b and CD62L) (18). Differences in methodology or gating strategy could explain these contradicting results. In general, we show that activation markers appear to increase upon ageing. This could possibly contribute to the low-grade inflammatory state that is known to be present in older adults (19).

We also show substantially decreased proportions of pDCs in the older group. This was in accordance with the findings of several other studies (20–22). Our study does however also show that this process is likely more dependent on chronological ageing rather than on the accumulation of deficits with age, as the HOC group was particularly healthy, reflected by their quality-of-life scores, and the lack of difference between frail and non-frail older people. This is in line with the prevailing explanation of the decrease in pDC proportions with age. pDCs are believed to originate from both myeloid and lymphoid progenitors (23). The most likely mechanism that underlies the decrease is a reduction in the output of pDCs from the bone marrow due to an unknown cause. The activation status of pDCs did not appear to be affected by age or by frailty in this study. Others however did show a decreasing capacity of pDCs to produce IFNα, which is their main effector function, which is dependent on intracellular TLRs, rather than TLR2/4 which we measured here (21,22). The decreased proportions of pDCs in the blood may underlie the impaired IFNα responses in older people, and may be one of the underlying reasons for impaired resilience against viral infections in older people.

Enhanced expression of CD40, which is generally very low on circulating monocytes, was found to be associated with frailty. We are, to the best of our knowledge, the first group to find an association with CD40 expression on monocytes and the frailty phenotype. Engagement of CD40 on monocytes was shown to induce production of pro-inflammatory cytokines, chemokines and metalloproteinases (24). Elevated CD40 levels have previously been found in patients with severe infections compared to healthy controls. However, the survival rate in these septic patients appeared to be lower in patients with lower CD40 levels, indicating that CD40 has important immunomodulating effects (25). Studies in bigger cohorts should reveal whether increased expression of CD40 in frail people impacts monocyte – T cell interactions and propagates inflammatory responses.

Our study has several strengths. We have included participants in three clearly defined groups based on age and frailty status. Frail and non-frail older adults could be compared because the groups were age matched. Frailty was primarily defined by the TFI, which may not be as widely used as the Frailty Index (8) Frailty Phenotype (7), but is still a validated and easily applicable tool to identify frailty. Moreover, the Frailty Phenotype was measured in a proportion of participants and showed, in coherence with the other quality-of-life questionnaires, clear differences between the frail and non-frail groups. Our flow cytometry panel was well-established for a clear and reliable identification of monocyte and DC subsets. MFI analyses allowed us to look at clear per-cell expression as opposed to the percentage of cells positive for activation markers. Unfortunately, we were unable to phenotype cDC1s further due to the small number of circulating cDC1 cells. Furthermore, the group size of n=15 is rather small and therefore results should be confirmed in other cohorts.

Our study is a descriptive study showing phenotypic changes of monocytes and DCs upon ageing and in frail older adults. Future studies should investigate the functional consequences of the increased activation status of monocytes and cDC2s upon ageing. Furthermore, the effect of increased CD40 expression by monocytes on the frailty phenotype should be investigated for functional consequences.

