## Supplementary data for "Effects of ageing and frailty on circulating monocyte and dendritic cell subsets"

Supplementary table 1: the 14 antibodies used in the flow cytometry panel

| **Marker** | **Fluorochrome** | **Company** | **Catalogue number** |
|---|---|---|---|
| CCR7 | PE | BD Biosciences | 552176 |
| CD1c | BUV395 | BD Biosciences | 742751 |
| CD11c | APC | BD Biosciences | 333144 |
| CD14 | Pacific Orange | Life Technologies | MHCD1430 |
| CD16 | BUV737 | BD Biosciences | 564434 |
| CD19 | AF-700 | Thermo Fisher Scientific | 56-0199-42 |
| CD40 | APC-Cy | BioLegend | 334224 |
| CD86 | BB515 | BD Biosciences | 564544 |
| CD141 | BB700 | BD Biosciences | 742245 |
| CD303 | BV785 | BioLegend | 354222 |
| HLA-DR | V450 | BD Biosciences | 655874 |
| PD-L1 | PE-Cy7 | BioLegend | 329715 |
| TLR2 | BV650 | BD Biosciences | 742769 |
| TLR4 | BV711 | BD Biosciences | 564404 |

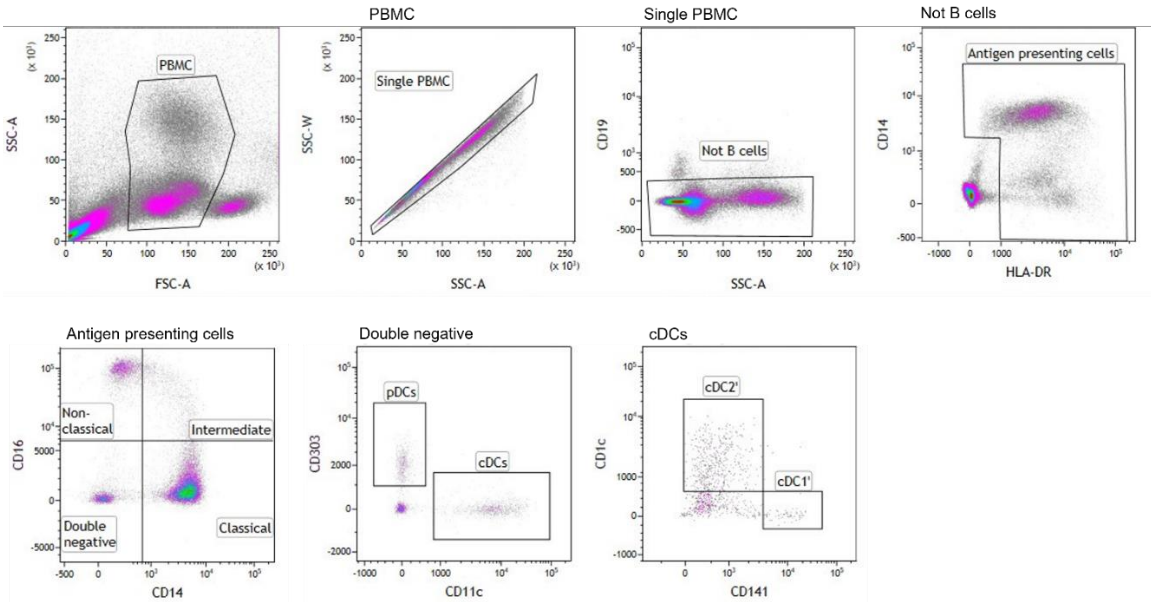


**Supplementary Figure 1: Gating strategy and marker expression in the flow cytometry experiments.** A: Single PBMCs were selected based on size and granularity. CD19 was used to gate out B cells. CD14 and HLA-DR was used to select all antigen presenting cells. Antigen presenting cells were further classified into non-classical, intermediate and classical monocytes based on CD16 and CD14 expression. CD16/CD14 double negative cells were divided into pDC based on CD303 and cDCs based on CD11c expression. cDC2 cells were CD1c positive and cDC1 cells were CD141 positive.

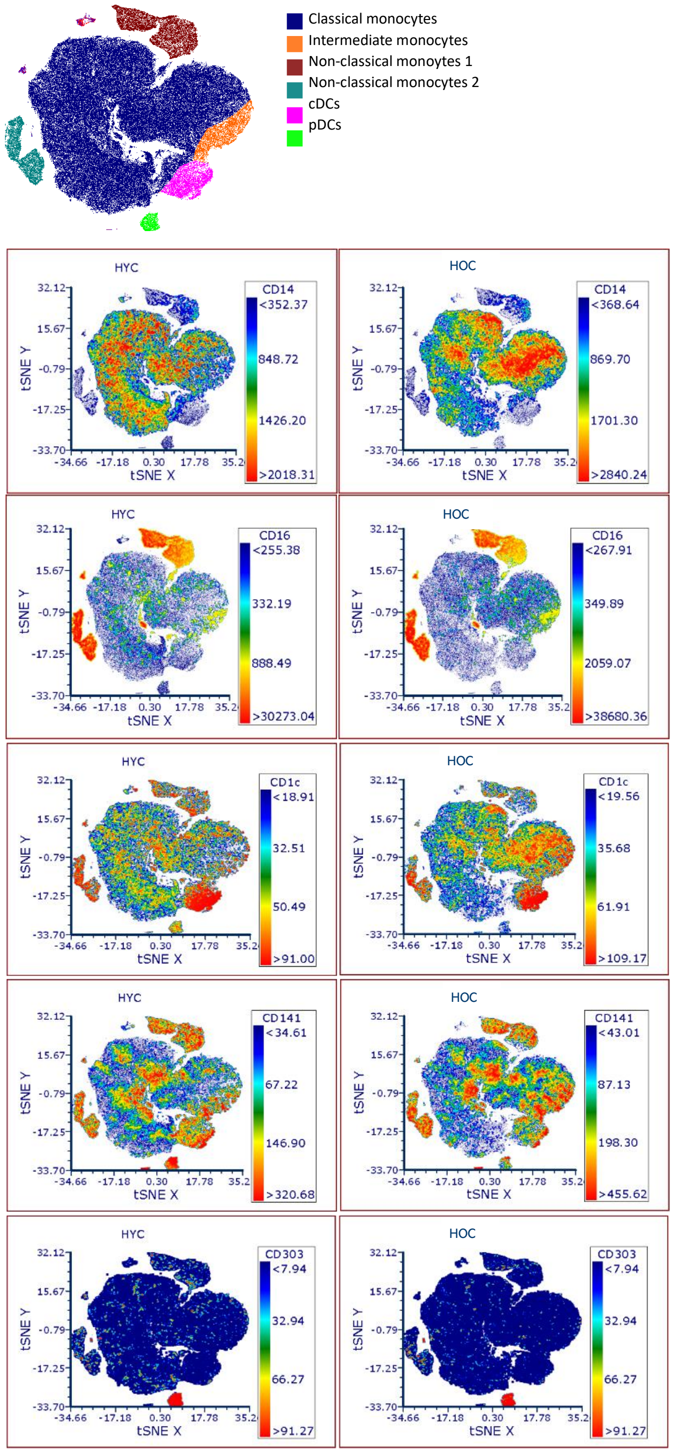


Supplementary figure 2: Expression of lineage markers CD14, CD16, CD1c, CD141 and CD303 in t-SNE plots of healthy young controls (HYC) and older healthy controls (HOC) for the identification of clusters with non-classical monocytes (CD14lowCD16+), intermediate monocytes (CD16+CD14+), classical monocytes (CD14+CD16-), conventional dendritic cells (CD141/CD1c+) and plasmacytoid dendritic cells (CD303+). t-SNE: t-distributed Stochastic Neighbor Embedding.

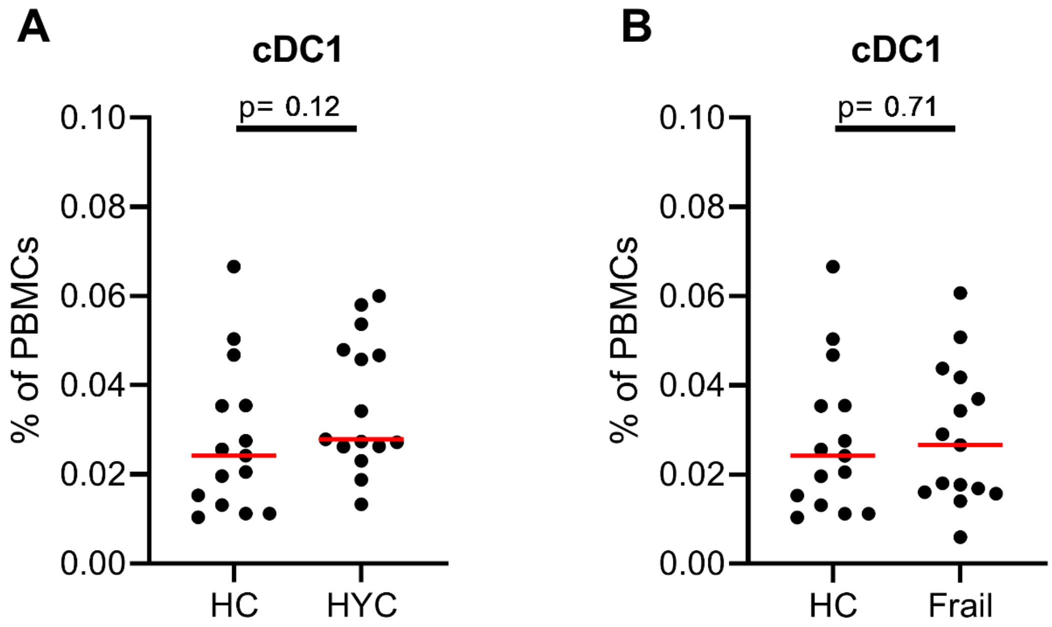


Supplementary figure 3: Proportion of cDC1 cells within total PBMCs. A: Proportion of cDC1 cells in older healthy controls (HOC) and young healthy controls (HYC). B: Proportion of cDC1 cells in older HOC and frail donors. The red line represents the median, and p-values of the Mann Whitney U test are shown in the graphs.

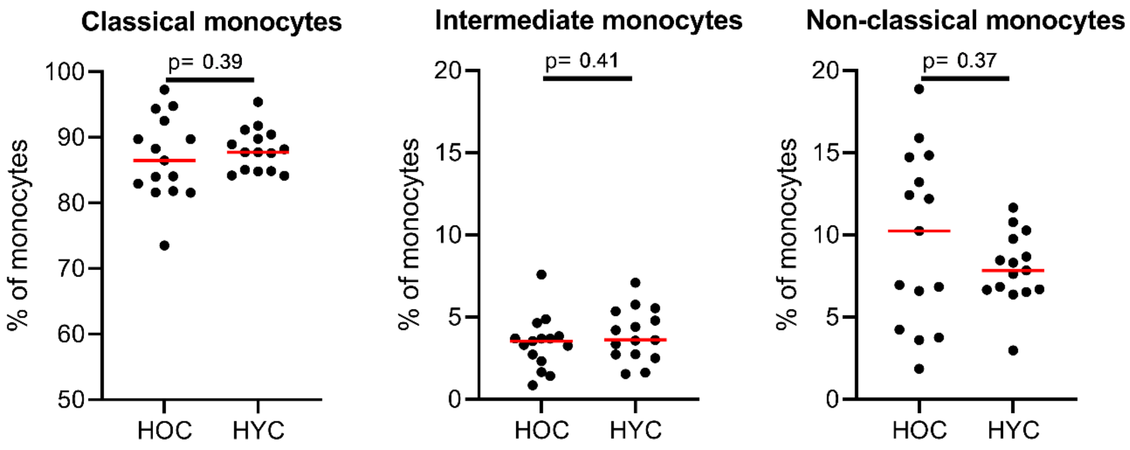


Supplementary figure 4: Proportion of monocyte subsets of total monocytes for the HOC and HYC groups. The red line represents the median, and p-values of the Mann Whitney U test are shown in the graphs. No significant differences between the HOC and HYC groups were found, indicating no evidence for a shift within monocyte subsets with age. HOC: healthy control, HYC: healthy young control

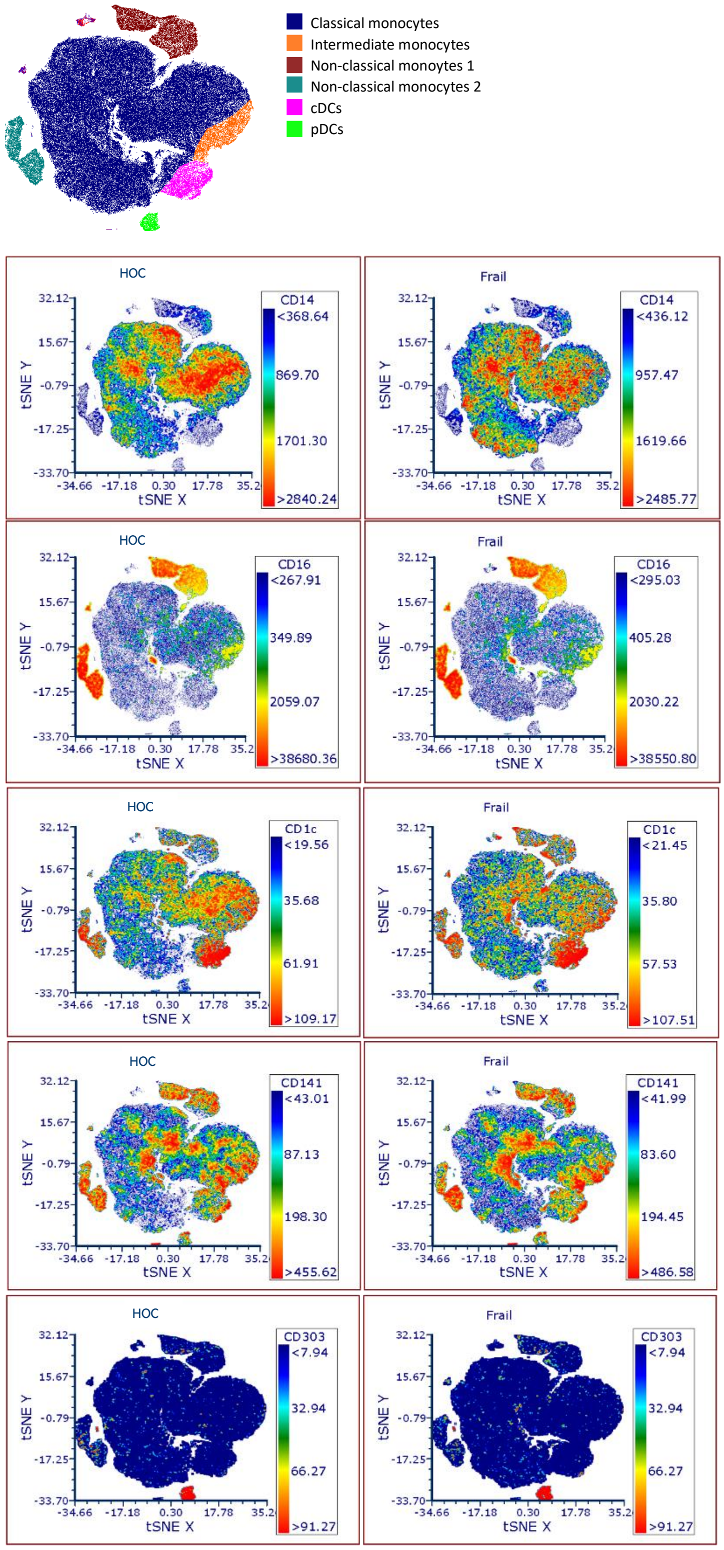


Supplementary figure 5: Expression of lineage markers CD14, CD16, CD1c, CD141 and CD303 in t-SNE plots of healthy older controls (HOC) and frail donors for the identification of clusters with non-classical monocytes (CD14lowCD16+), intermediate monocytes (CD16+CD14+), classical monocytes (CD14+CD16-), conventional dendritic cells (CD141/CD1c+) and plasmacytoid dendritic cells (CD303+). t-SNE: t-distributed Stochastic Neighbor Embedding.

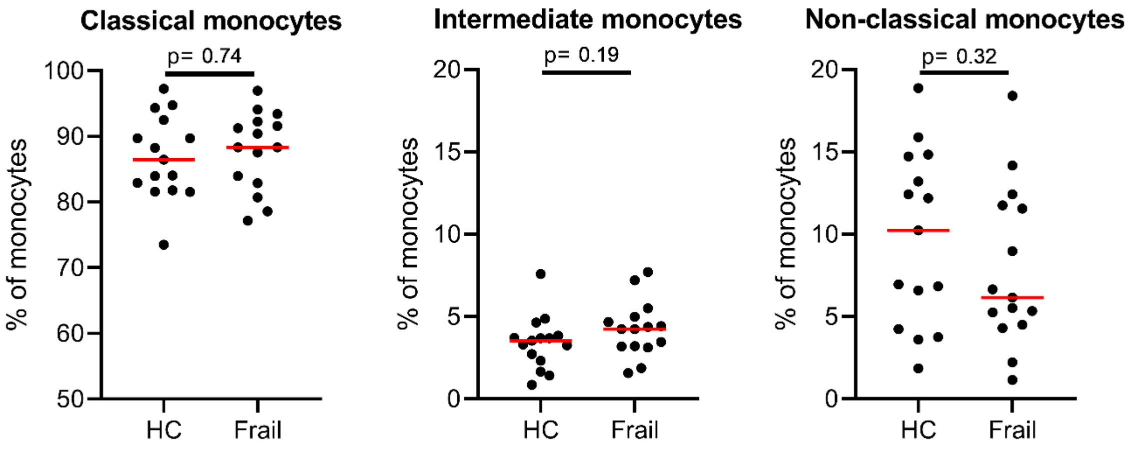


Supplementary figure 6: Proportion of monocyte subsets of total monocytes for the HOC and Frail groups. The red line represents the median, and p-values of the Mann Whitney U test are shown in the graphs. No significant differences between the HOC and Frail groups were found, indicating that in frail older people there is no evidence for a shift within monocyte subsets. HOC: healthy control.
